# Excitatory-Inhibitory Balance within EEG Microstates and Resting-state fMRI Networks: Assessed via Simultaneous PET-MR-EEG Imaging

**DOI:** 10.1101/2020.05.22.109413

**Authors:** R Rajkumar, C Régio Brambilla, T Veselinović, J Bierbrier, C Wyss, S Ramkiran, L Orth, M Lang, E Rota Kops, J Mauler, J Scheins, B Neumaier, J Ermert, H Herzog, KJ Langen, F Binkofski, C Lerche, NJ Shah, I Neuner

## Abstract

The symbiosis of neuronal activities and glucose energy metabolism is reflected in the generation of functional magnetic resonance imaging (fMRI) and electroencephalography (EEG) signals. However, their association with the balance between neuronal excitation and inhibition (E/I-B), which is closely related to the activities of glutamate and GABA (γ -aminobutyric acid) and the receptor availability (RA) of GABA_A_ and mGluR5, remains unexplored. This study investigates these associations during the resting state (RS) condition using simultaneously recorded PET/MR/EEG (trimodal) data. Glucose metabolism and neuroreceptor binding availability (non-displaceable binding potential (BP_ND_)) of GABA_A_ and mGluR5 were found to be significantly higher and closely linked within core resting-state networks (RSNs). The neuronal generators of EEG microstates and the fMRI measures were most tightly associated with the BP_ND_ of GABA_A_ relative to mGluR5 BP_ND_ and the glucose metabolism, emphasising a predominance of inhibitory processes within in the core RSNs at rest.

## Introduction

It is generally acknowledged that the functional architecture of the brain is governed by fundamental principles such as functional segregation and integration(Zeki and Shipp, 1988). Functional integration (both intrinsic and induced) in the brain is directed by the balance between neuronal excitation and inhibition and is closely related to activity in the brain’s main excitatory and inhibitory neurotransmitters, glutamate and GABA (γ -aminobutyric acid), respectively(Duncan, Wiebking and Northoff, 2014). This excitatory/inhibitory balance (E/I-B) has been shown to play an important role in several crucial mental processes(Carcea and Froemke, 2013). It is postulated that this balance is the underlying mechanism influencing both intrinsic and stimulus-induced activity across brain regions and may also play a role in the constitution of the higher cognitive functions, such as self and consciousness(Duncan, Wiebking and Northoff, 2014). Conversely, disturbances in the E/I-B may lead to functional network alterations and have been shown to be underlying factors in some psychiatric diseases(Allen *et al*., 2019). Thus, a profound understanding of the neurobiological basis of the E/I-B remains a major scientific challenge and a deeper comprehension of it would offer greater insight into psychiatric diseases.

The E/I-B has been extensively investigated at the cellular level of neural activity but remains less well described at the more complex levels where it becomes interspersed with various aspects of brain function. Details relating to the mechanistic aspects of the E/I-B are assessable using different neuroimaging methods, with each method providing information pertaining to a somewhat different aspect of neural activity. Functional magnetic resonance imaging (fMRI) records the haemodynamic response - the blood oxygenation level-dependent (BOLD) contrast - which is likely to be driven by balanced proportional changes in the excitation and inhibition of neurons(Logothetis, 2008). Multichannel electroencephalography (EEG) reflects voltage changes resulting from the synchronous firing of groups of neurons in the brain and thus mainly exposes the neuroelectric activity at the synapse. Positron emission tomography (PET) uses specific radioligands to visualise and quantify a variety of metabolic and physiological processes *in vivo*.

Alone, each of these techniques provides valuable insight into the examined processes in terms of its own frame of reference. However, each technique is also constrained by its own limitations. In recent years, the development of a combined approach – simultaneous trimodal PET-MR-EEG imaging(N.J. Shah *et al*., 2017) - has enabled the measurement of different aspects of the same process as it occurs, under the same physiological and psychological conditions, shedding valuable light on the relationship between them. Thus, this innovative method is highly suitable to elucidate the coupling between neuronal activities, energy consumption, oscillations and the E/I-B.

To date, our investigations using simultaneous trimodal MR-PET-EEG imaging have mainly focused on the default mode network (DMN), as this is the first described and one of the most prominent resting-state networks (RSN), containing brain regions critical for several cognitive functions. Using the trimodal approach, it was possible to demonstrate a significantly higher glucose metabolism (quantified by a 2- [^18^F]fluoro-2-desoxy-D-glucose PET (FDG-PET)) in the DMN compared to the whole-brain non-DMN grey matter (GM) during resting state (RS)(N.J. Shah *et al*., 2017). Further, the neuronal activation within the DMN (as assessed with fMRI) was positively correlated with the metabolic activity (assessed as the mean standard uptake value of FDG). Electrical source localisation of EEG signals also showed a unique frequency range pattern.

In another investigation, a more detailed clarification of the associations between the EEG signals and the ongoing glucose energy metabolism during rest was sought by specifically focussing on EEG microstates (Rajkumar *et al*., 2018). The spatial distribution of active neuronal sites that contribute to electrical activity in the brain can be revealed using EEG scalp topographies. Such topographies are found to be quasi-stable for periods of about 80-120 ms and are known as EEG microstates(Lehmann, 1990). RS EEG studies have consistently identified four microstate maps (topographies)(Koenig *et al*., 2002), each being generated by diverse neuronal assemblies. Their locations can be estimated using source localisation techniques, such as the standard low-resolution brain electrotomography (sLORETA)(Pascual-Marqui, 2002). The transition of microstate maps may be regarded as a sequential activation of various neuronal assemblies. Van De Ville *et al*. have indicated the microstates as being “atoms of thought” (the shortest constituting elements of cognition)(Van De Ville *et al*., 2010). A small number of studies have investigated the relationship between EEG microstates and RS-fMRI using simultaneously recorded RS-EEG and fMRI, and have highlighted the four typical microstates likely to represent the resting-state networks (RSNs) usually identified in RS-fMRI studies(Britz, Van De Ville and Michel, 2010; Van De Ville *et al*., 2010). With our own trimodal approach, we demonstrated tight associations between fMRI metrics, FDG-PET standard uptake values and the single microstates, indicating a functional relationship between cortical hubs, connectivity parameters and the glucose metabolism(Rajkumar *et al*., 2018).

However, our understanding of these complex inter-connections in the context of the E/I-B is very sparse. As the EEG microstates from large-scale neural networks are a result of synaptic electrical activity due to the excitatory and inhibitory actions of the neurotransmitters, we hypothesise that the neuronal generators of EEG microstates are tightly associated not only with the functional connectivity measures and glucose metabolism but also with the availability of the neuroreceptors relevant for the E/I-B. With this in mind, we have chosen the following two receptors as representatives of the E/I-B for our investigation: the excitatory G-protein-coupled metabotropic glutamate receptor type 5 (mGluR5) and the inhibitory, ligand-gated, ion-specific GABA_A_ receptor. For their quantification within the PET part of the trimodal MR-PET-EEG imaging, two selective radioligands were used to determine receptor availability (RA): [^11^C]ABP688 (ABP), for mGluR5(Treyer *et al*., 2007) and [^11^C]Flumazenil (FMZ) for GABA_A_ receptors (Hansen *et al*., 1991). Further, 2-[^18^F]-fluoro-2-desoxy-D-glucose (FDG) was used to study glucose metabolism. During each of these three separate investigations, fMRI and EEG data were assessed simultaneously with identical protocols. The data obtained were analysed jointly in an explorative approach in order to address the following questions:

1. To what extent does the E/I-B (expressed as the RA of mGluR5 and GABA_A_) and the glucose metabolism measured via FDG-PET determine the activity of the neuroelectric generators of microstates as estimated by sLORETA during RS?
2. Is the previously shown higher metabolic activity within the DMN during the RS associated with a higher RA of the mGluR5 and GABA_A_ as an expression of the higher energy demand to maintain the E/I-B in the RS?
3. In view of the tight association of the neuronal activity (regardless of whether excitatory and inhibitory) with the glucose metabolism, is there a link between the RA of the mGluR5 and GABA_A_ and the glucose metabolism measured via FDG-PET?
4. In light of the known modulatory effect of the excitatory and inhibitory neurotransmission on the BOLD signal(Lecrux *et al*., 2011; Just and Sonnay, 2017), is there an association between the RS functional connectivity and the RA of mGluR5 and GABAA within the RSNs?

## Results

The simultaneous trimodal data acquisition was successfully completed in 29 subjects. Each modality is discussed in detail below.

### PET measures

The PET data were reconstructed and parametric images representing BP_ND_ and SUV were calculated as described in the methods section. The calculated PET measures are shown in Fig. 1.

**Fig. 1:**
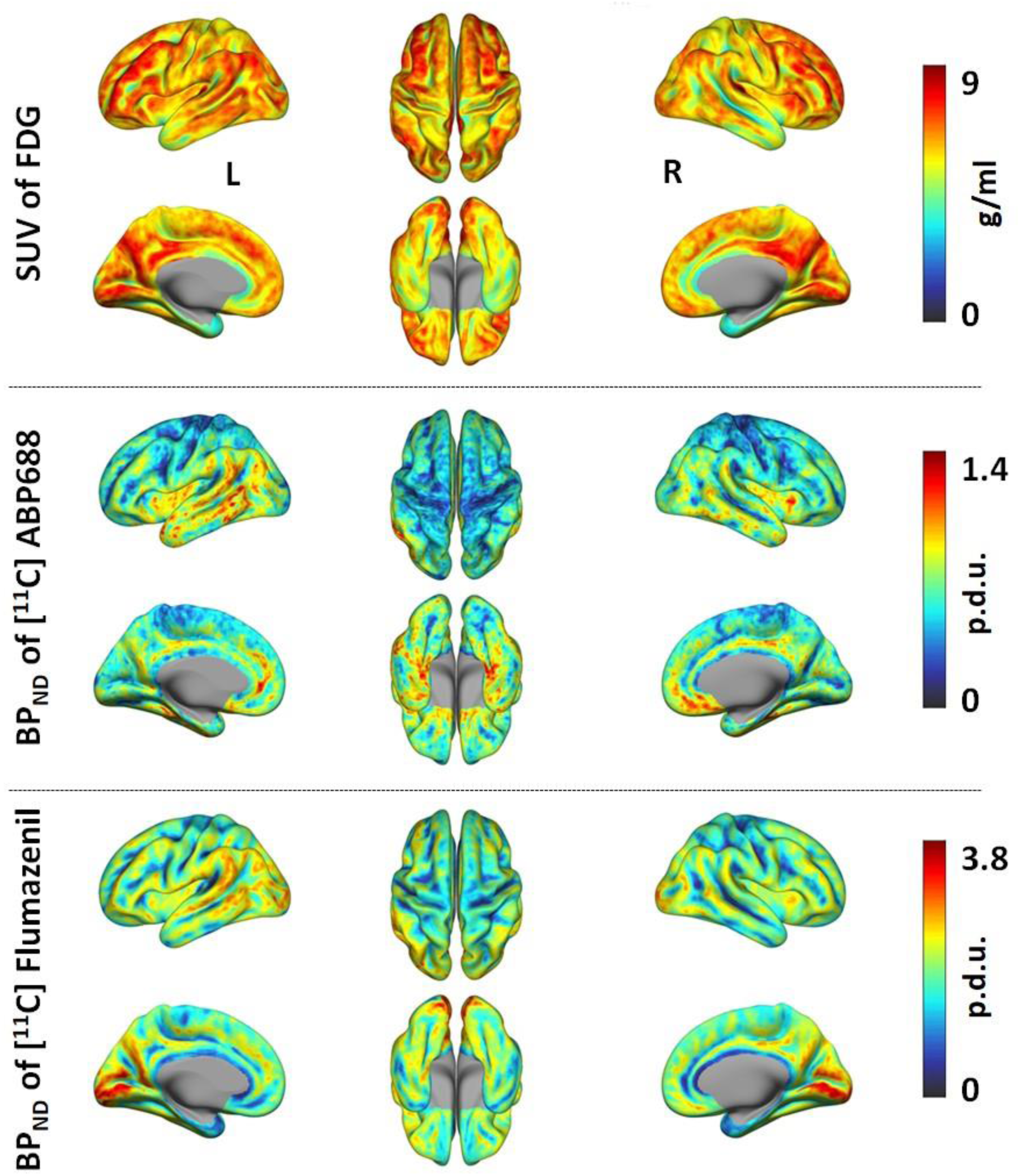
PET parametric images. The 3D images representing the SUV of the FDG tracer are shown in the top row; the BP_ND_ of ABP and FMZ are shown in the middle and bottom rows. The SUV images show glucose consumption in g/ml, the BPND images show the distribution of the neuroreceptor availabilities for mGluR5 (middle row) and GABA_A_ (bottom row) receptors in procedure defined unit (p.d.u.). The calculated PET parametric images, averaged across all subjects in each study, are presented. The left, superior and right views are shown in the upper rows and left medial, inferior and right medial views are shown in the lower rows of each parametric images.

### EEG microstates

The artefacts in the EEG data were removed and the microstates were calculated. The 3 dimensional (3D) cortical current density distribution for each microstate was calculated using the sLORETA method. The calculated microstates and the 3D cortical current density distribution of each microstate are shown in Fig. 2.

**Fig. 2:**
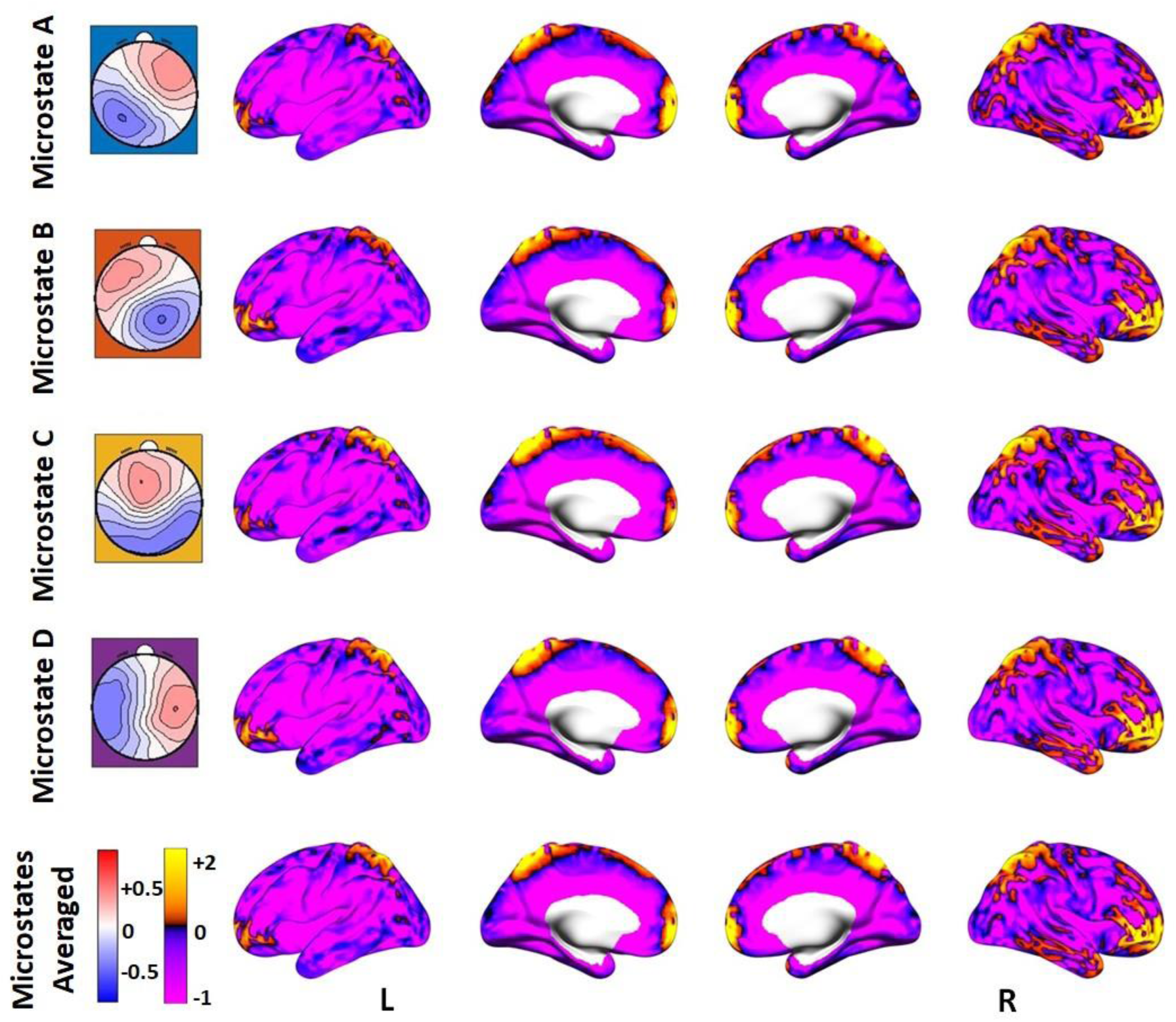
EEG microstates. The topographic maps of the EEG microstates from the mean template of all subjects (first column) and corresponding Z-value standardised 3D source-localised images (averaged across all subjects) showing cortical current density distribution. The positively coloured regions in the 3D images depict the regions of neuronal generators of EEG microstates. The bottom row images show the average image of the four microstates. The left, left medial, right medial and right views are shown in each row of the 3D source-localised images.

### *Association* between receptor availability and EEG microstates

The Pearson linear correlation coefficients computed between the PET and source-localised microstate measures showed significant, low positive correlations (p<0.01) (Fig. 3) with corrections for multiple comparison. The correlation coefficients (r) are summarised in Supplementary Table. 1.

**Fig. 3:**
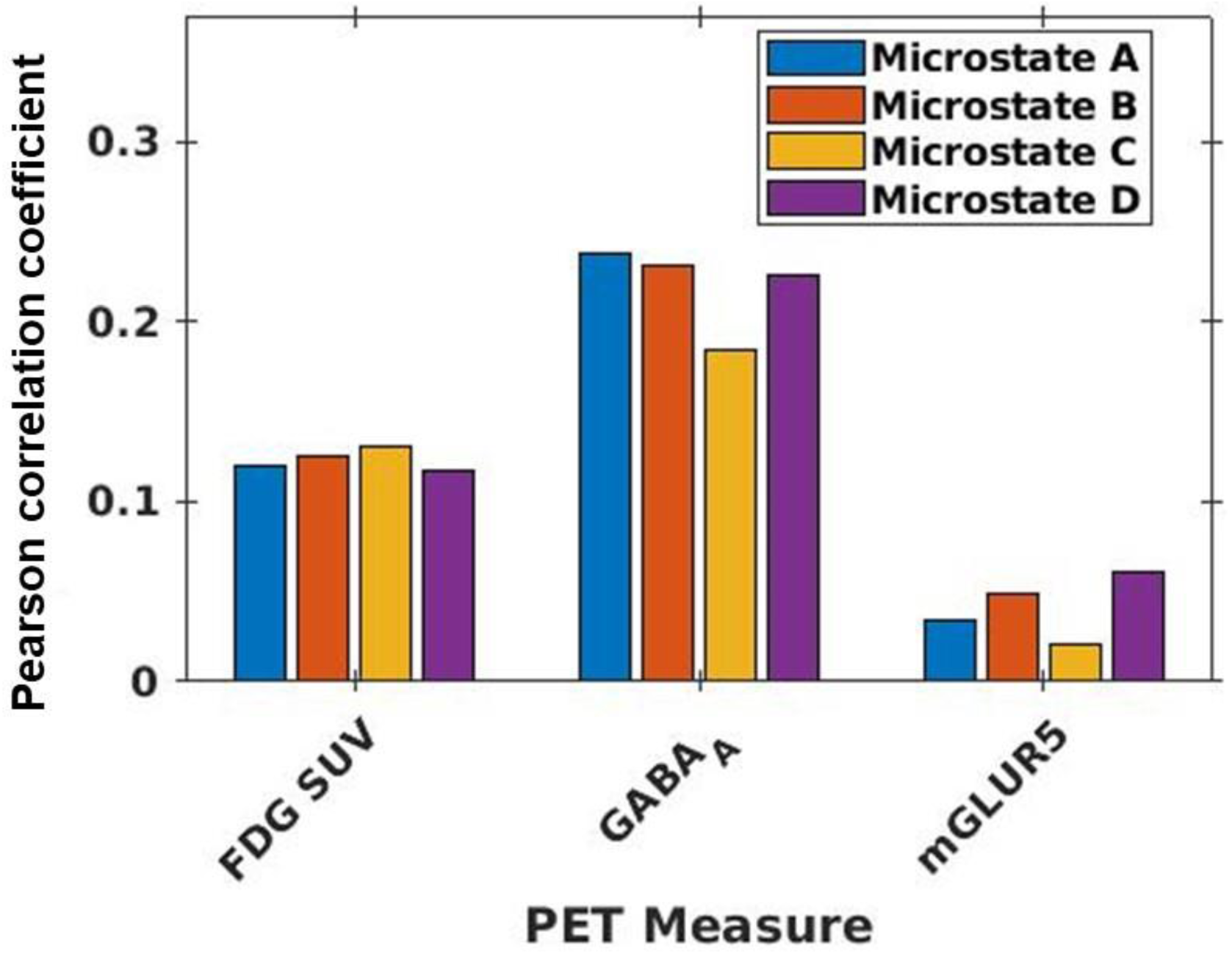
Bar chart showing the Pearson correlation values between PET and microstates. The correlation values (p <0.01) were plotted computed between averaged values within microstate masks from FDG-PET SUV, receptor availabilities of GABA_A_, and mGluR5 with the source-localised microstate maps from EEG.

### RS-fMRI measures

The fMRI data were successfully pre-processed. The RS-fMRI measures (ReHo, DC and fALFF) were calculated for each subject as described in the methods section. The calculated RS-fMRI measures averaged across all the 29 subjects are shown in Fig. 4.

**Fig. 4:**
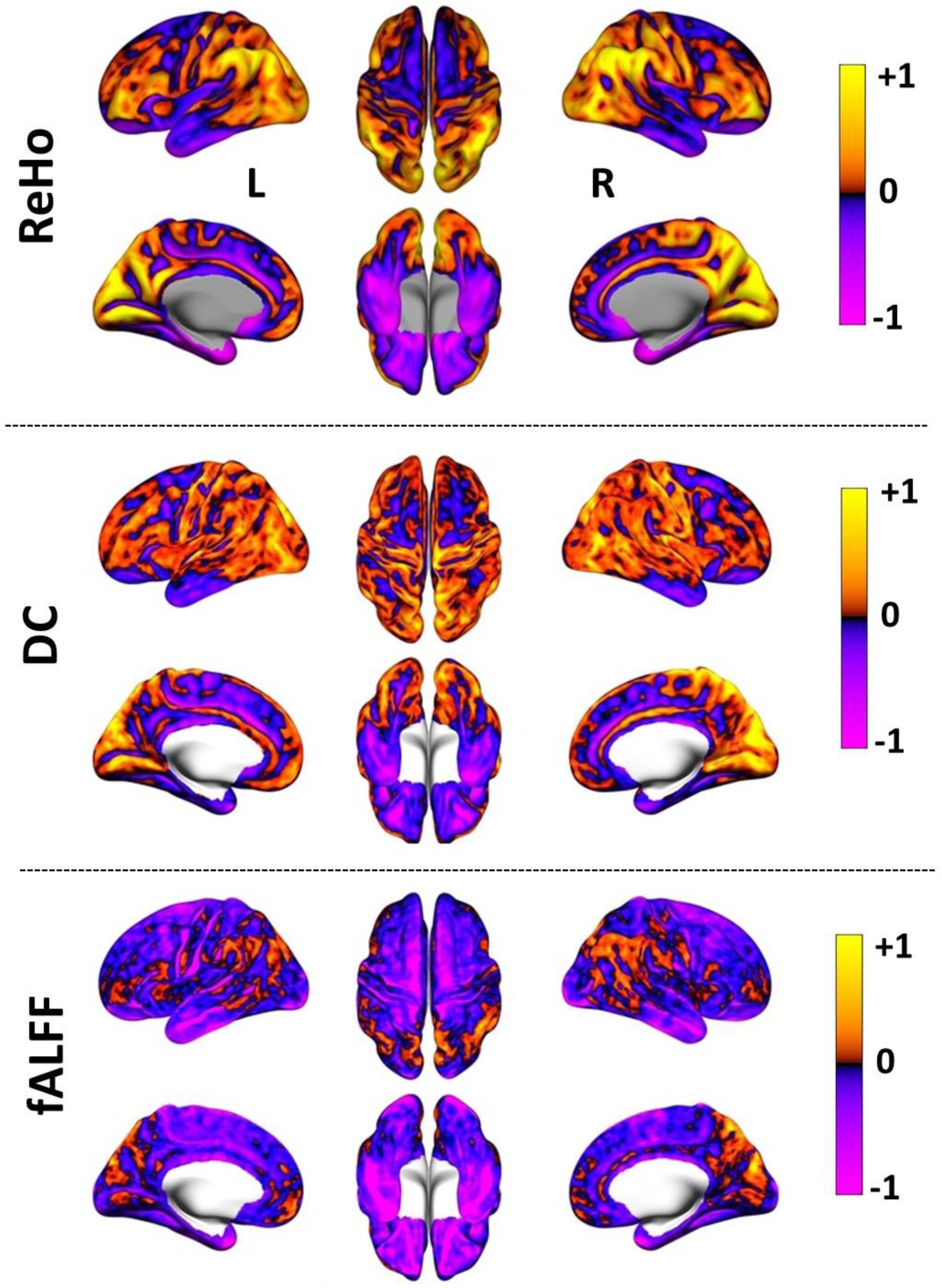
RS-fMRI measures. Z-value standardised measures 3D images representing regional homogeneity (ReHo - top row), degree centrality (DC - middle row) and fractional amplitude of low frequency fluctuations (fALFF - bottom row) RS-fMRI measures averaged across all subjects are shown. The positively coloured regions in the 3D images show higher short-range functional connectivity in ReHo, long-range functional connectivity in DC and BOLD signal fluctuations in fALFF. The left, superior and right views are shown in the upper rows and left medial, inferior and right medial views are shown in the lower rows of each RS-fMRI measure.

### Receptor availabilities

The Dunn’s test showed that the RA of GABA_A_ and mGluR5 in the core RSNs were significantly (p<0.05) higher compared to GM or SMN (Fig. 5) for the within-PET measure analysis.

**Fig. 5:**
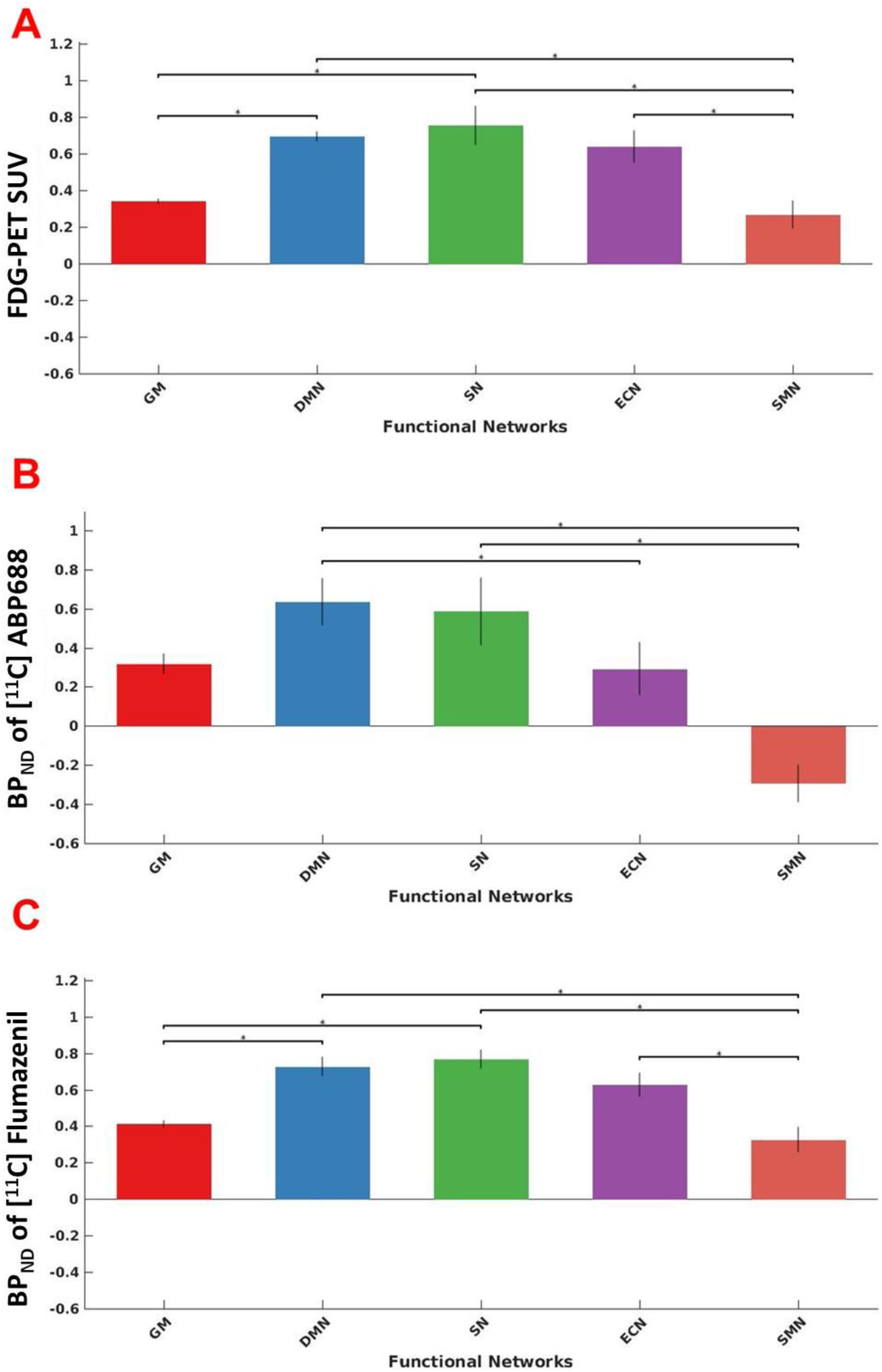
Bar plots showing FDG-PET SUV (A) and the distribution of neuroreceptor availability of mGLUR5 (B) and GABA_A_ (C) in whole-brain grey matter (GM), default mode network (DMN), salience network (SN), executive control network (ECN) and sensory motor network (SMN). The error bars show the standard deviations. The horizontal markers with an asterisk symbol show the significant differences between regions based on the Dunn’s test.

### Association between receptor availability and glucose metabolism

The one-sample KS test showed the extracted voxel values from the averaged PET, fMRI and EEG measures to follow a normal distribution in all cases. The Pearson linear correlation coefficients (r) computed between the BP_ND_ values and FDG-PET SUV showed significant positive correlations (p<0.01) within GM as well as the core RSN regions. The correlation results after family wise error correction are summarised in the bar plot (Fig. 6).

**Fig. 6:**
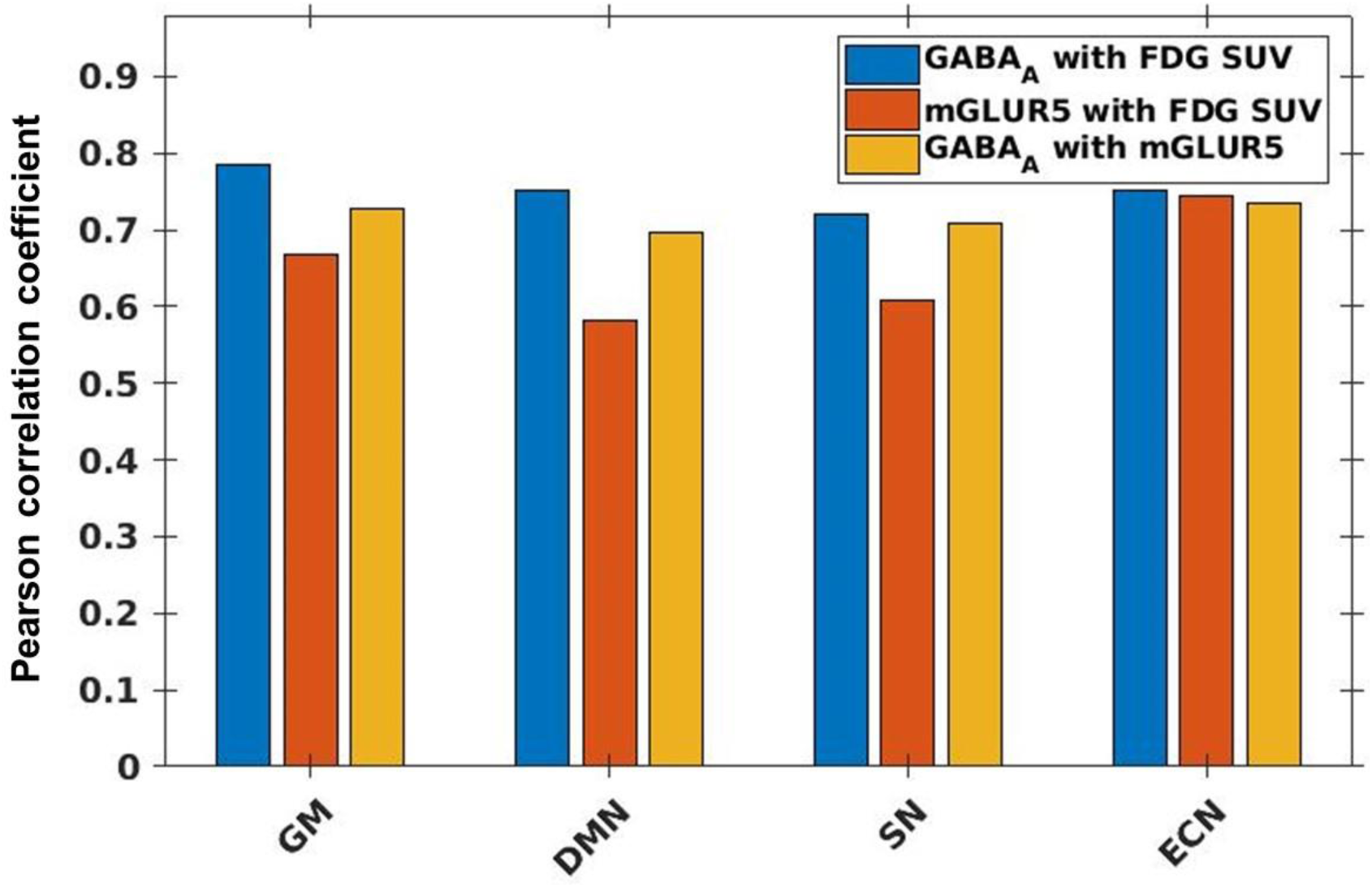
Bar plot showing Pearson correlation coefficient values calculated between receptor availability of GABA_A_ with FDG-PET SUV (blue bars), receptor availability of mGluR5 with receptor availability of GABA_A_ (red bars), and receptor availability of GABA_A_ with receptor availability of mGluR5 (yellow bars). The correlation values are shown for the whole-brain grey matter region (GM), the default mode network (DMN), the salience network (SN) and the executive control network (ECN).

### Association between receptor availability and RS-fMRI measures

The Pearson linear correlation coefficients computed between the PET and fMRI measures showed significant positive correlations (p<0.01) within GM as well as the core RSNs (Fig. 7) with corrections for FWER. The correlation coefficients (r) are summarised in Supplementary Table. 2.

**Fig. 7:**
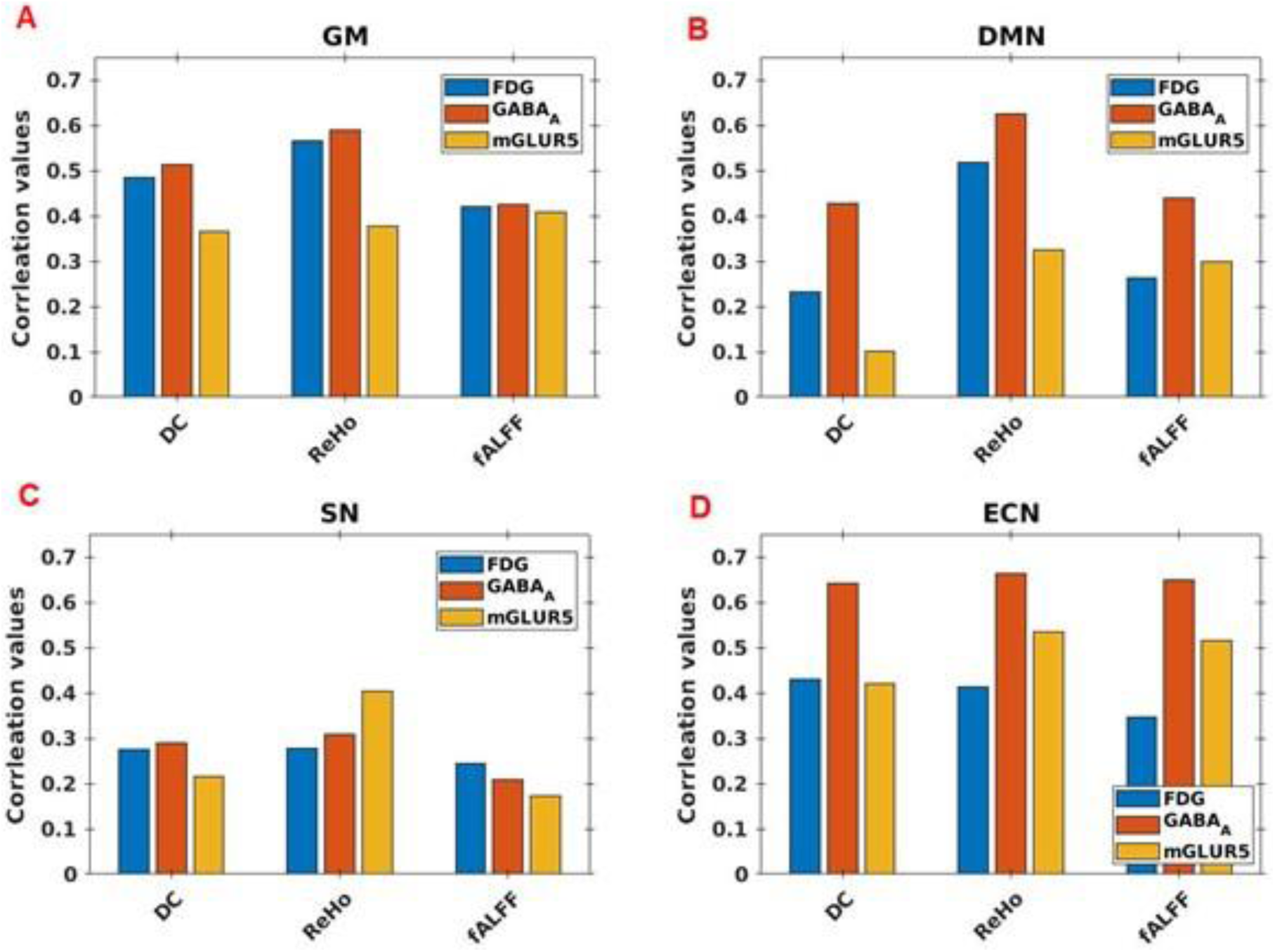
Bar chart showing the Pearson correlation values between PET and fMRI measures in the whole-brain grey matter (GM)), the default mode network (DMN), the salience network (SN) and the executive control network (ECN). Correlations between PET and fMRI regional homogeneity (ReHo), degree centrality (DC) and fractional amplitude of low-frequency fluctuation (fALFF) measures are shown.

## Discussion

The E/I-B is crucial for the maintenance of the brain’s functional integration. In this study, we examined the neurobiological correlates involved in E/I-B within resting-state fMRI networks using simultaneously recorded PET-MR-EEG trimodal imaging data. This unique technology enabled elucidation of the coupling between neuronal activities, energy consumption, neuronal generators of EEG microstates and the availability of the mGluR5 and the GABA_A_ neuroreceptors - fundamental for excitatory and inhibitory neurotransmission. Our study demonstrates both the feasibility and advantages of measuring trimodal data simultaneously using different radioligands as well as illustrating a new approach to studying associations between neuronal activities through various measures derived from combined trimodal data. For the first time, insight into the neurobiological foundation of the E/I-B (given as the quantification of the availability of the excitatory and inhibitory receptors as well as the glucose metabolism intensity) has been gained alongside the functional connectivity measures and EEG microstates during the RS.

### EEG microstates

The source-localised microstate maps shown in Fig. 2 demonstrate the cortical current density distribution (neuronal generators) of each microstate, with all microstates showing the highest current density distribution in the posterior cingulate cortex. Additional current density distributions were seen in the anterior cingulate, and in the left and right occipital as well as the parietal cortices. Having averaged the microstates, each microstate was found to represent a fragmented version of the DMN (Fig. 2). According to the cortical current density distribution map of the microstates, the DMN regions were restricted to the cortical areas only usually identified via fMRI or PET. In line with previous observations(Pascual-Marqui *et al*., 2014), the source-localised microstates proved the DMN regions, observed by means of fMRI or PET imaging, to be a low-pass filtered version of the microstate dynamics.

### GABA_A_ and mGluR5 receptor availability and their associations with EEG microstates

The results depicted in Fig.1, show the averaged BP_ND_ images of FMZ and ABP, corresponding to the distribution of neuroreceptor availability of GABA_A_ and mGluR5, respectively. These corroborate the notion that GABA_A_ receptors are widely distributed throughout the human brain(Waldvogel, Baer and Faull, 2010), whereas mGluR5 receptors are mostly distributed in the anterior cingulate cortex, the medial temporal lobe, the putamen and the caudate nucleus(Ametamey *et al*., 2007).

In addition, the results of this study show a significantly higher RA of GABA_A_ in the DMN and SN compared to whole-brain GM as well as in all of the core RSNs compared to the non-core RSNs (SMN) (Fig. 5 (C)). Thus, even though GABA_A_ is widely distributed in the brain, the significantly higher RA in the RSNs suggests greater inhibitory action, particularly during resting conditions.

With respect to the RA of mGluR5, while it was not found to be higher in any of the core RSNs compared to whole-brain GM (Fig. 5 (B)), it was found to be significantly higher in the two core RSNs DMN and SN compared to the SMN (Fig. 5 (B)). Thus, the higher RA of both GABA_A_ and mGluR5 receptors in the core RSNs compared to SMN during RS indicates that these networks, in particular, require a stable equilibrium between excitation and inhibitory processes compared to the SMN.

Our results pertaining to the link between the cortical generators of EEG microstates with the RA of GABA_A_ receptors show comparatively higher correlation values than with the FDG-PET SUV and with the RA of mGluR5 receptors (Fig. 3). This is in support of other EEG studies which have shown the involvement of GABA_A_ receptors in the beta oscillations (13-28 Hz)(Porjesz *et al*., 2002). Furthermore, studies using MRS have also demonstrated an association between GABA concentration and EEG signals in the beta band in the sensorimotor cortex(Baumgarten *et al*., 2016), auditory gating in schizophrenia(Rowland *et al*., 2013), and in the working memory load processing capacity in the dorsolateral prefrontal cortex(Yoon, Grandelis and Maddock, 2016). As microstates represent a fragmented low-pass filtered version of the DMN and considering that GABA_A_ receptors mainly mediate synaptic inhibition, our results confirm, from a different perspective, the importance of the inhibitory processes mediated by GABA_A_ receptors in the DMN during rest.

Despite the higher RA of mGluR5 seen in the DMN (Fig. 5 (B)), the neuronal generators of EEG microstates did not show substantial correlation with the RA of mGluR5 (Fig. 3). mGluR5 receptors are reported to regulate various mechanisms implicated in neurogenesis and synaptic maintenance(Piers *et al*., 2012) as well as in long-lasting forms of synaptic plasticity, including long-term depression (LTD) and long-term potentiation (LTP)(Kullmann and Lamsa, 2008). Further, the mGluR5 receptors share a structural and functional link with the NMDA-R and jointly modulate glutamatergic signalling(Newell and Matosin, 2014). In studies combining EEG and PET measures, an association between the mGluR5 RA and EEG delta activity could be demonstrated(Holst *et al*., 2017). Keeping in mind the proven association between the glutamate concentration and EEG oscillations in the gamma band(Lally *et al*., 2014), the negligible association between mGluR5 receptors and the neuronal generators of EEG microstates accessed in the low-frequency range in our study indicate that the excitation of neurons during RS is likely mediated by other glutamate receptors in the 2-20 Hz frequency range.

### Glucose metabolism and its association with the availability of mGlu5 and GABAA receptors

Glucose is the main energy substrate of the adult brain. Although the brain represents only 2% of the body mass, it requires about 20% of the oxygen and 25% of the glucose consumed by the human body(Bélanger, Allaman and Magistretti, 2011). Thereby, the basal metabolism (required for vital functions of neurons) accounts for 30% of the energy consumed by the brain with the remaining 70% being accounted for by spontaneous activity(Tomasi, Wang and Volkow, 2013).

In our study, we replicate a significantly higher FDG uptake (corresponding to a higher glucose metabolism) in the core RSNs - DMN and the SN-compared to the SMN and the whole-brain GM regions in the resting state, as already previously shown in other studies (Tomasi, Wang and Volkow, 2013; N.J. Shah *et al*., 2017). Previous investigations have also demonstrated that, relative to this very high rate of ongoing energy consumption in the RS, the additional energy consumption associated with changes in brain activity is remarkably small, often less than 5% of the baseline level of activity(Raichle and Mintun, 2006). Our results indicate that the main proportion of the energy consumption during the RS is directed to the highly active DMN and SN. A task-related deactivation of the DMN(Anticevic *et al*., 2012) may be permissive for an energy redirection to other task-related networks. The high energy demand of the SN during rest may be associated with the continuous state of readiness required within the scope of its responsibility for detecting and filtering information necessary to maintain goal-directed behaviour by shifting attention between external and internal processes(Menon and Uddin, 2010).

Regarding the question relating to the associations between the glucose metabolism measured via FDG-SUV and the RA of the mGluR5 and GABA_A_, strong voxel-level correlations were observed with both neuroreceptor types in all examined core RSNs, as well as in the whole-brain GM (Fig. 6).

These results correspond with the established knowledge that glutamate cycling consumes a large proportion of the total energy utilized by the brain(Attwell and Laughlin, 2001) and that 60-80% of the total energy consumption in the brain during RS is devoted to glutamate cycling(Sibson N. R. *et al*., 1997). Glutamate is the principal neurotransmitter responsible for the majority of excitatory synaptic activity. However, it exhibits very complex signalling and functional diversity, which is decoded by several different receptor types assigned to the ionotropic (iGluR) and the metabotropic glutamate receptor family (mGluR). In this case, mGluR5 belongs to the group I mGluRs family and is usually localised postsynaptically. In contrast with group II and group III mGluRs, which are often localised on presynaptic terminals or preterminal axons where they inhibit neurotransmitter release, activation of group I mGluRs often leads to cell depolarisation and increases in neuronal excitability(Niswender and Conn, 2010). Thus, mGluR5 receptor may be considered as a suitable representative of excitatory glutamatergic neurotransmission and is tightly coupled with glucose metabolism.

We further observed a strong correlation between the GABA_A_ receptors, which are widely distributed in the cerebral cortex, and the FDG-PET SUV in the core RSNs and whole-brain GM (Fig. 6). GABA_A_ receptors are pentameric ligand-gated ion channels mainly composed of two α, two β and a single γ subunit, which are selected from a diverse pool of 19 different subunit types(Sigel and Steinmann, 2012) and are responsible for most fast synaptic inhibition in the mammalian brain. They also play a crucial role at extra-synaptic sites by mediating tonic inhibition in the brain(Jacob, Moss and Jurd, 2008). The tight association between the glucose homeostasis and the GABAergic neurotransmission has been demonstrated in some previous PET studies(Kim *et al*., 2014). Here we confirm these findings by demonstrating a tight interrelation between the GABA_A_ RA and glucose metabolism, measured directly, in vivo.

Finally, high correlation values were found between mGluR5 and GABA_A_ in all core RSNs as well as in the remaining GM (Fig. 6). Considering that these are the two most important representatives of the receptors required for excitatory and inhibitory processes in the brain, our results indicate a neurobiological basis of the E/I-B by the means of the innovative trimodal approach.

### Glucose metabolism and its association with EEG microstates

Several previous studies have demonstrated tight associations between glucose concentration and metabolic rate and EEG signal changes(Blaabjerg and Juhl, 2016). A direct association between the glucose metabolism and the neuronal generators of EEG-microstates has, to the best of our knowledge, not been investigated before now. In our study, rather surprisingly, a low association was observed between the FDG-SUV and the neuronal generators of the four EEG-microstates (Supplementary Table 1). Given that the synaptic currents responsible for neuronal firing (recorded via EEG during RS) display a high energy demand, mainly provided by glucose, a stronger correlation was expected between the cortical generators of EEG microstates and the glucose consumption indicated by the SUV of FDG. One possible explanation for the rather weak correlation values revealed here may be due to the different observational periods: the FDG-PET SUV values are showing the FDG uptake averaged over about 6 minutes while the microstates lasted only a few milliseconds.

### Associations of the fMRI measures with glucose metabolism and mGluR5 and GABA_A_ RA

The fMRI techniques utilise the blood oxygenation level-dependent (BOLD) contrast to map the brain’s functional integration. The fMRI signal in the DMN is influenced by the energy demands associated with synaptic currents and action potential propagation(Attwell and Laughlin, 2001). In the absence of external stimuli, the RS-fMRI signal partially represents the metabolic processes but also fluctuations of spontaneous neuronal activity(Fukunaga *et al*., 2008). Thus, examining the relationship between the SUV using FDG-PET and fMRI measures will likely facilitate an accurate interpretation of the BOLD signals.

In this study, the RS-fMRI measures (DC, ReHo, and fALFF) showed a significant voxel-level correlation with FDG-PET SUV measures in whole-brain GM (Fig. 7 (A)), which is in agreement with previously reported results(Aiello *et al*., 2015). This indicates a high interrelation between metabolic processes and spontaneous neuronal activity - the high energy demand of the brain seems to be utilised for high functional connectivity (as revealed in DC and ReHo) as well as for the BOLD fluctuations (as revealed by fALFF).

However, the correlation between the fMRI measures and the SUV from FDG-PET is weaker in the core RSNs compared to GM (Fig. 7 (B-D)). This may be explained due to the high energy efficiency of the RSNs(Tomasi, Wang and Volkow, 2013), which, despite having more functional connections, possibly use less energy. Further, our results indicate a stronger association of FDG-PET SUV with ReHo than with DC in the DMN (Fig. 7 (B)). This may be on account of the more short-range, energy-consuming connections (ReHo) within the DMN or a higher energy-efficiency of the longer-range connections (DC). Such difference was not observed within the SN and the ECN, where within both networks the correlation of the FDG-PET SUV values with ReHo and DC was similarly strong. This points towards a unique property of the DMN relating to energetic efficiency.

To date, the association between the fMRI measures and the glutamatergic and GABAergic neurotransmission has primarily been investigated using the single-voxel magnetic resonance spectroscopy (MRS) technique. In task conditions, depending on the brain regions and cognitive processes involved, GABA levels have shown both positive and negative correlations with the BOLD signal(Northoff *et al*., 2007; Levar *et al*., 2017). Arrubla *et al*. (2014) have found GABA concentration in the posterior cingulate cortex (PCC) to be inversely correlated with connectivity within the DMN during RS(Arrubla *et al*., 2014). Furthermore, both resting-state and task-based fMRI studies(Enzi *et al*., 2012; Duncan, Wiebking and Northoff, 2014) have also shown associations with glutamate concentration levels. However, the drawback of single-voxel MRS is that the neurotransmitter’s concentration from a single voxel (typically a 3 cm isotropic cube) is used to investigate connectivity within RSNs or the whole brain. In simultaneous MR-PET imaging, this drawback is surmounted by calculating three-dimensional parametric images of the whole brain (about 3mm resolution), which enables region-of-interest (ROI) analysis.

Our results show disparate association patterns between the fMRI measurements and the GABA_A_ and mGluR5 RA for the three core RSNs and GM, as depicted in Fig.7. Regarding the associations between the long-range connectivity measure (DC) and the excitatory and inhibitory RA (mGluR5 and GABA_A_), the most prominent difference was observed in the DMN. Here we found an almost fourfold higher association with the GABA_A_ RA than with the mGluR5 RA. A similarly directed, although albeit lower relation, was also observed for the parameters ReHo and fALFF, indicating a predominant role of the inhibitory GABAergic neurotransmission within the E/I-B in the DMN.

Interestingly, a higher association between the short-range measure ReHo and the mGluR5 RA was only observed in the SN. The SN is considered to be a filtration and amplification network that is responsible for evaluating specific external/internal stimuli and assigning the relevance of stimuli for goal-directed behaviour(Menon, 2015). In this specific operating range, the SN may request a higher capacity of excitatory (compared to the inhibitory) receptors and connections than within other RS networks.

### Summary

In summary, the results presented above contribute to a better understanding of the questions initially raised in the introduction and reveal the following pertinent conclusions:

1. The main, relatively high significant correlation of all four EEG microstates was observed with the RA of GABA_A_ receptors, followed by considerably lower correlations with the FDG-PET SUV and correlations with the RA of mGluR5 receptors to a very low extent. Considering that microstates represent a fragmented low-pass filtered version of the DMN, these results confirm the importance of the inhibitory processes mediated by GABA_A_ receptors in the electrophysiological activity of the DMN during resting states. The considerably lower association between the EEG microstates and the FDG-PET SUV appears somewhat surprising and may be related to methodology. The negligible association between mGluR5 receptors and the neuronal generators of EEG microstates accessed in the low-frequency range indicate that the excitatory contribution in the E/I-B may be mediated by other glutamate receptors than mGluR5.
2. A significantly higher distribution of GABA_A_ and mGluR5 receptors was observed in both the DMN as well as in the SN compared to the non-core RSNs (SMN). However, compared to the whole GM, only the RA of GABA_A_ was significantly higher in these networks. This finding once again supports a leading role of the inhibitory processes during the RS. Furthermore, the RA of mGluR5 and GABA_A_ correlated significantly with each other in all core RSNs and in the remaining GM, indicating the general importance of E/I-B during the RS.
3. The RA of the mGluR5 and GABA_A_ receptors both showed strong voxel-level correlations the glucose metabolism measured via FDG-SUV in all examined core RSNs as well as in the whole-brain GM. This finding indicates that the high energy demand during the RS may be mainly directed to the maintenance of the E/I-B.
4. Each of the observed core RSNs and GM showed specific association patterns between the different fMRI measure’s and the GABA_A_ and mGluR5 RA. Thus, the relative proportion of excitatory and inhibitory processes within the E/I-B seems to differ between the single RSNs and is most likely to be dependent on their main functions.

### Limitations and outlook

Several methodological limitations should be kept in mind when interpreting our results. One of the main limitations of the study is the relatively small sample size, a fact hampering a broader generalization of our findings. Furthermore, the investigation combines data from three separate studies, each using just one PET tracer in a separate subject group. However, conducting all three examinations in the same group is not conceivable due to the complexity of the trimodal data acquisition paradigms as well as due to the ethical and legal implications relating to the repeated radiation exposure from PET imaging. Moreover, the radiation safety authority in the authors’ country of research has imposed a 10-year restriction on the injection of PET tracers for the second time in healthy individuals for research purposes. To reduce the implications of this limitation, we included very well-matched groups in the three separate investigations.

Receptor availability was only examined in mGluR5 and GABA_A_ receptors, which represent a small fraction of the excitatory and inhibitory neuroreceptors. Investigating all neuroreceptors, or at least the most prominent ones, would have helped to delineate the mechanisms behind the neurovascular and neuroelectric signals in greater detail. Nevertheless, the two chosen receptor types appear to be the most representative of the receptors required for excitatory and inhibitory neurotransmission.

Further, the radioligands used for the quantification of the mGluR5 and GABA_A_ RA differ in their receptor binding properties, but appear to show similar high receptor affinity (for ABP: dissociation constant (K_D_)= 1.7 ± 0.2 nmol/L(Ametamey *et al*., 2006); for FMZ K_D_= 0.7 ± 0.26 nM(Pandey *et al*., 1997)).

These limitations notwithstanding, the results presented here give new insight into the neurobiological foundation of the E/I-B (given as the quantification of the availability of the excitatory and inhibitory receptors as well as the glucose metabolism intensity) in terms of functional connectivity measures and EEG microstates during the RS. Further research involving healthy individuals as well as patients with mental illness will likely shed valuable light towards a better understanding of the neurobiological alterations underlying specific diseases.

## Methods

The trimodal data presented in this study were acquired using three different PET – radiotracers: [^11^C]ABP688, [^11^C]Flumazenil and [^18^F]FDG. For readability, the datasets are hereafter referred to as ABP, FMZ and FDG data, based on the radiotracer used in the study.

### Trimodal data acquisition

The simultaneous trimodal data were acquired using a 3T hybrid MR-BrainPET scanner system (Siemens, Erlangen, Germany)(Herzog *et al*., 2011) equipped with an MR-compatible EEG system (Brain Products, Gilching, Germany). As shown in a previous study, the EEG electrodes used in this system do not cause any noticeable effect on PET images(Rajkumar *et al*., 2017). The study was approved by the Ethics Committee of the Medical Faculty of RWTHAachen University and the German Federal Office for Radiation Protection (*Bundesamt für Strahlenschutz*). All methods were performed according to the relevant guidelines and regulations. The study was conducted in accordance with the Declaration of Helsinki and prior written consent was obtained from all volunteers. Only the trimodal FDG data presented in this study were taken from a previously published work (N. J. Shah *et al*., 2017). Only subjects (or their first degree relatives) with no history of neuropsychiatric disorders as assessed via Mini International Neuropsychiatry Interview (MINI)(Sheehan *et al*., 1998) were included in each study. In addition, only male subjects were included due to the effects of the menstrual cycle on variations in neurotransmitter levels(Harada *et al*., 2011). The data and results presented in this manuscript will be available upon request via the corresponding author.

The data acquisition protocol for each modality is described below.

#### PET data acquisition

Radiosynthesis of [^11^C]ABP688 and [^11^C]Flumazenil were performed according to literature(Canales-Candela, Riss and Aigbirhio, 2012; Elmenhorst *et al*., 2016).The PET radioligand was injected as bolus in the FDG study and as bolus plus infusion in the ABP and FMZ studies. The radiotracer was injected via an intravenous (IV) line while the subject was lying in the 3T hybrid MR-BrainPET scanner. Details relating to the number of subjects, age, and injected activity with K_Bol_, for each study are given in Table 1. PET data acquisition in list mode started immediately following injection of the tracer. In ABP and FMZ study, the eyes closed RS-fMRI and EEG data were simultaneously recorded from the time the radiotracer reached equilibrium in the brain. In the FDG study where no equilibrium is reached, the resting state data was recorded exactly from 50 minutes after radiotracer injection in all subjects. During RS data acquisition, subjects were instructed to close their eyes and not to fall asleep. The simultaneous acquisition of RS data lasted for approximately 6 minutes.

**Table 1:** Subject and injected activity details for each study

| ***Study*** | ***Number of subjects*** | ***Mean age***<br>***in years*** | ***Injected activity***<br>***in MBq*** |
| --- | --- | --- | --- |
| **ABP** | 9 (Males) | $23 \pm 3$ | $407 \pm 56$<br>($K_{\text{Bol}} = 61.79 \text{ min}$) |
| **FMZ** | 10 (Males) | $26 \pm 3.72$ | $411 \pm 18$<br>($K_{\text{Bol}} = 46.22 \text{ min}$) |
| **FDG** | 10 (Males) | $28 \pm 4$ | $200 \pm 30$ |

Table 1: Subject and injected activity details for each study
| ***Study*** | ***Number of subjects*** | ***Mean age***<br>***in years*** | ***Injected activity***<br>***in MBq*** |
| --- | --- | --- | --- |
| **ABP** | 9 (Males) | $23 \pm 3$ | $407 \pm 56$<br>($K_{\text{Bol}} = 61.79 \text{ min}$) |
| **FMZ** | 10 (Males) | $26 \pm 3.72$ | $411 \pm 18$<br>($K_{\text{Bol}} = 46.22 \text{ min}$) |
| **FDG** | 10 (Males) | $28 \pm 4$ | $200 \pm 30$ |

The PET data were iteratively reconstructed (3D OP-OSEM, 32 iterations and 2 subsets) into 2-minute frames. Attenuation correction was performed based on an MR- and template-based procedure(Rota Kops *et al*., 2015). In addition, the PET data reconstruction incorporated corrections for random and scattered coincidences, dead time, radioactive decay, and pile up. The reconstructed image frames comprised 256 × 256 pixels and 153 slices (voxel size 1.25 mm^3^, isotropic).

#### MR data acquisition

In all investigations, the anatomical and RS-fMRI images were acquired using magnetisation-prepared rapid acquisition gradient-echo (MP-RAGE) and T2*-weighted echo planar imaging (EPI) sequences, respectively. The anatomical images were acquired before the RS data acquisition during the same session. The MR imaging parameters considered in each sequence are given below,

- MP-RAGE sequence: repetition time (TR) = 2250 ms, echo time (TE) = 3.03 ms, field-of-view (FOV) 256 × 256 × 176 mm^3^, voxel size = 1×1×1 mm^3^, flip angle (FA) = 9°, 176 sagittal slices and a GRAPPA acceleration factor of 2 with 70 auto-calibration signal lines.
- EPI sequence (for FDG and FMZ studies): TR = 2200 ms, TE = 30 ms, FOV = 200 × 200 × 108 mm^3^, voxel size = 3.125 × 3.125 × 3.0 mm^3^, FA = 80°, number of slices = 36, number of volumes = 169
- EPI sequence (for ABP study): TR = 2000 ms, TE = 30 ms, FOV = 192 × 192 × 108 mm^3^, voxel size = 3 × 3 × 3.0 mm^3^, FA = 80°, number of slices = 33, number of volumes = 180

#### EEG data acquisition

Prior to the trimodal acquisition, the MR-compatible EEG cap (BrainCap MR, EasyCap GmbH, Herrsching, Germany) was placed on the subject’s head and an electrolyte gel (ABRALYT 2000, EASYCAP GmbH, Herrsching, Germany) was applied between the EEG electrodes and the scalp to reduce impedance. The impedance of all recording electrodes was kept below 10 kΩ. A 32-channel EEG cap was used in the FDG study and a 64-channel EEG cap was used in the ABP and FMZ studies. The EEG electrodes were positioned according to the 10 - 20 system in both caps. The electrocardiogram (ECG) signals were recorded using an additional electrode. The ECG electrode was connected to the distal part of the trapezius muscle. The RS EEG data (eyes closed) and ECG were recorded using the Brain Vision Recorder (Brain Products, Gilching, Germany) while the subject lay in the scanner. The subjects were instructed to stay calm and relaxed and not fall asleep during the RS measurements. The RS EEG data were recorded at a sampling rate of 5 kHz and bandwidth between 0.016 and 250 Hz. The helium pump of the MR system was switched off during trimodal RS data acquisition in order to avoid the effects of vibration on the MR and EEG system as well as on the subjects.

### Data analysis

#### PET

The PET images were analysed using the PMOD (version 3.5) software package. The reconstructed PET images were corrected for motion using a PET image-based motion correction method (Scheins *et al*., 2019). Following motion correction, the PET images were smoothed with a Gaussian kernel size of 2.5 mm along all three directions. Only the three PET image frames corresponding to the RS measurement were considered for further analysis.

The parametric images representing the non-displaceable binding potentials (BP_ND_)(Mintun *et al*., 1984) in each subject were calculated from the reconstructed ABP and FMZ PET data. The three PET image frames were averaged (time weighted). The BP_ND_ was calculated as (C_T_ - C_REF_)/ C_REF_, where C_T_ denotes the activity concentration in an area of high neuroreceptor density, and C_REF_ denotes activity concentration in a non-specific binding region (region with very low receptor density). The cerebellum(Akkus *et al*., 2017) and pons(Odano *et al*., 2009) were chosen as C_REF_ regions for ABP and FMZ, respectively. The C_REF_ regions were defined using a three-dimensional maximum probability atlas(Hammers *et al*., 2003) on the MNI space co-registered anatomical images (MP-RAGE).

The parametric images representing the SUV were calculated from FDG-PET data using the averaged PET image frame during RS. The total activity injected (MBq) and the body weight (kg) of the subject were accounted for when calculating the SUV. The final unit of SUV is g/ml.

#### fMRI

Several methods can be employed to analyse the RS-fMRI data (Lee, Smyser and Shimony, 2013), computing various RS-fMRI measures with respect to the connectivity and strength of low-frequency BOLD fluctuations. In this study, RS-fMRI measures such as regional homogeneity (ReHo)(Zang *et al*., 2004), degree centrality (DC)(Buckner *et al*., 2009), and fractional amplitude of low-frequency fluctuations (fALFF)(Zou *et al*., 2008) are considered. The chosen RS-fMRI measures are distinct, data driven and do not require prior definition of seed voxels/regions. While the ReHo measure only considers the neighbouring voxels, thus depicting short-range functional connectivity, the DC measure takes into account all voxels and is considered a long-range functional connectivity measure. The slow BOLD signal fluctuations are characterised by the fALFF measure. Of the currently identifiable RSNs, the DMN, the salience network (SN) and the executive control network (ECN)(Menon, 2011) are considered to be the core neurocognitive RSNs and, as such, form the basis of analysis in this study. Furthermore, for the purposes of RA comparison within the RSNs, the sensory motor network (SMN) (a non-core RSN) is also considered as a control region.

The RS-fMRI measures (ReHo, DC, fALFF) were calculated using MATLAB based software packages, SPM12 (http://www.fil.ion.ucl.ac.uk/spm/) and DPABI(Yan *et al*., 2016). The pre-processing steps included the removal of the first 10 volumes of the total acquisition, slice-timing correction with respect to the middle slice of the functional image, realignment, and nuisance covariates regression (NCR). The covariates for NCR included head motion parameters(Friston *et al*., 1996), whole-brain white matter (WM) and cerebrospinal fluid (CSF) mean signals, and the constant, linear and quadratic trends in the BOLD signals. Following the pre-processing steps, RS-fMRI measures were calculated in the subject’s native image space. The temporal filtering of BOLD fMRI signals between 0.01 and 0.08 Hz was performed before DC and ReHo calculation. The ReHo measure was calculated using Kendall’s coefficient of concordance (KCC)(Stuart, 1956) as the homogeneity measure over 27 neighbouring voxels(Li *et al*., 2012). The DC measure was calculated with the Pearson correlation threshold of 0.25 (p = 0.001). The fALFF measure was calculated within the BOLD low-frequency range between 0.01 and 0.1 Hz.

#### EEG

The EEG data were processed using the MATLAB-based open-source software package EEGLAB version 13(Delorme and Makeig, 2004) and analysed in three steps. First, the data pre-processing was performed to remove artefacts, then the microstates were calculated, and in the final step the microstates were source-localised. Following is a detailed explanation of the EEG processing steps.

##### EEG data pre-processing

The recorded EEG data were imported to EEGLAB and down-sampled to 2048 Hz. When recorded alongside fMRI, EEG data are heavily contaminated by gradient artefacts (GA) due to the switching of the gradient magnets in the MR system. Here, the GA were corrected using the FASTR tool(Niazy *et al*., 2005) implemented in EEGLAB. The data were then downsampled to 1000 Hz, filtered between 2 and 20 Hz using a Hamming windowed sinc finite impulse response (FIR) filter implemented in the EEGLAB function *eegfiltnew*, and re-referenced to the average reference. Bad channels were identified and removed based on their correlation to neighbouring channels using the EEGLAB function *clean_rawdata*. In order to remove non-stationary artefacts, such as head movement, artefact subspace reconstruction (ASR)(Mullen *et al*., 2015) was performed with the recommended parameters(Chang *et al*., 2018). Following ASR, the data were again re-referenced to the average reference. The pulsatile flow of blood due to the pumping action of the heart causes ballistocardiogram (BCG) artefacts in the EEG data. An adaptive optimal basis set algorithm (aOBS)(Marino *et al*., 2018), which accounts for the key shortcomings of the commonly used optimal basis set (OBS) algorithm(Niazy *et al*., 2005), was used to detect and remove these artefacts from the EEG signal. An automatic blind source separation algorithm was used to remove ocular artefacts(Go’mez-Herrerol *et al*., 2006). To remove further residual artefacts, independent component analysis (ICA)(Bell and Sejnowski, 1995) was performed and the multiple artefact rejection algorithm (MARA)(Winkler, Haufe and Tangermann, 2011) was used to classify and remove artefactual components. Further BCG-related components were identified and removed with the projection onto independent components (PROJIC) algorithm(Abreu *et al*., 2016). Finally, the first 30 seconds of the RS-EEG data were rejected, as was the case in the fMRI analysis.

##### Microstate Calculation

The microstates in the artefact-corrected EEG data were analysed using the Microstate plugin (http://www.thomaskoenig.ch/index.php/software/microstates-in-eeglab) implementation in EEGLAB. Here, the global brain activity is represented in the global field power (GFP) of the multichannel EEG data(Lehmann and Skrandies, 1980), with the EEG topographies being considered to be stable around the maxima of the GFP (GFP peaks) (Koenig *et al*., 2002). Thus, the GFP was first calculated for each subject, and the EEG topographic maps were then plotted for GFP peaks. All EEG topographic maps marked as GFP peaks were extracted and submitted for spatial clustering, which was performed using a modified atomised and agglomerate hierarchical clustering (AAHC) algorithm(Tibshirani and Walther, 2005). The AAHC algorithm was set to identify the dominant EEG topographic maps for each subject. In the next step, the dominant EEG topographic maps across all subjects in each study (group template) were identified from the single-subject EEG topographic maps. The group template was also identified using the modified AAHC algorithm, which resulted in a group template map for each study. Thus, an ABP EEG-MS group template, an FMZ EEG-MS group template, and an FDG EEG-MS group template were obtained. The group-template topographic maps from each study were then automatically sorted into microstates A, B, C and D using the template maps provided in the Microstate plugin, as reported in a previous RS microstate study(Koenig *et al*., 2002). Finally, the EEG topographic maps identified for each subject were also sorted into microstates A, B, C and D, using the sorted group-template maps from each study.

In order to reduce the computation time, the EEG data were again downsampled to 125 Hz. Following the microstate calculation, sorting, and downsampling, the EEG data corresponding to each microstate were epoched separately as EEG-MS A, EEG-MS B, EEG-MS C, and EEG-MS D for every individual subject in each study. Within epoched EEG-MS, the GFP peaks were again identified and the microstates that lasted less than 50ms were rejected. By considering the identified GFP peak as the centre, 50ms of EEG data were epoched from both sides of GFP peak for each microstate in EEG-MS. The multichannel EEG data lasting 50ms surrounding the GFP peak were chosen, as previously mentioned, to ensure the stability of the microstates around the GFP peaks. All the 50ms epoched EEG data within each EEG-MSs were averaged for each subject separately. This resulted in an averaged 50ms multichannel EEG data around the GFP peak for each EEG-MS A, B, C, and D from every subject. For example, in the ABP study, a 50ms averaged EEG data epoch was created for each microstate, EEG-MS A, B, C, and D, with respect to every individual subject. The averaged 50ms EEG data epochs were then used for the source localisation, which is detailed in the next step.

##### Source localisation

The 50ms EEG data epochs were exported to the LORETA-KEY software (http://www.uzh.ch/keyinst/loreta.htm) to solve the inverse problem (source localisation). In a first step, the EEG electrode positions in the MNI152 template were determined(Jurcak, Tsuzuki and Dan, 2007) and a 3D cortical current density distribution from the 50ms EEG data epochs was calculated using sLORETA(Pascual-Marqui, 2002). In order to solve the inverse problem, sLORETA computations were made in the realistic head model(Fuchs *et al*., 2002) using the Montreal Neurological Institute template (MNI152)(Mazziotta *et al*., 2001). The 3D solution space in sLORETA was restricted to the cortical grey matter (GM). The inverse solutions showing the cortical current density was averaged for each microstate, resulting in a 3D cortical current density distribution for each microstate from every subject.

#### Intermodality comparison

In order to perform intermodality comparisons, the calculated measures from PET, fMRI and EEG were co-registered to the MNI152 (2×2×2 mm^3^) standard space. Additionally, the co-registered images were linearly standardised into Z-values(Aiello *et al*., 2015) for each subject. Z-value standardisation was calculated by subtracting the mean whole-brain voxel value from each voxel and then dividing the difference by the standard deviation of the whole brain. The Z-value standardised measures were then smoothed with a Gaussian kernel size of 3 mm along all three directions.

#### RSNs definition

Masks of the RSNs were obtained from an atlas of 90 functional regions of interest (fROIs)(Shirer *et al*., 2012). In order to further restrict the analysis to the GM regions, only the voxels within the GM region of the RSNs (which show more than 50% probability of being GM) were considered (GM correction). A GM tissue-segmented MNI152 (2×2×2 mm^3^) template (GM mask) was used for GM correction. A whole-brain GM mask was created by considering only the voxels that showed more than 50% probability of being GM from the tissue-segmented MNI152 template (Supp. Fig. 1). Given that different subjects were used in each study (FDG, FMZ and ABP), a subject-level comparison of PET values within each calculated PET measure is not possible. Thus, all the SUV and BPND images were averaged, yielding an averaged PET measure image. Similarly, for fMRI measures, and in order to improve the signal-to-noise ratio, averaged fMRI measure images were calculated for DC, ReHo and fALFF for each study. Subsequently, all voxel values within the GM-corrected RSNs and GM masks were extracted from the previously calculated subject-level PET, averaged PET and averaged fMRI measures.

Since the 3D solution space in sLORETA was restricted to only the cortical GM, a separate mask for each microstate (MS mask) was created considering only the voxels in the 3D solution space. The microstate maps across subjects were averaged for each study. Finally, all voxel values within the MS masks were extracted from the previously averaged PET, fMRI and EEG measures.

#### Statistics

All statistical analyses were performed using the MATLAB (Release 2017, The MathWorks, Inc., Natick, United States) software package. The mean values of SUV and BP_ND_ were calculated from each subject’s PET image (FDG, FMZ and ABP) for all of the considered regions (GM, DMN, SN, ECN, SMN and MS masks). In order to compare the differences in each PET measure within the considered regions, a stepdown Dunn’s test for non-parametric multiple comparisons(Dunn, 1961) was performed with a significance level of 5%. Dunn’s test was performed using a MATLAB implementation(Cardillo, 2006). A one-sample Kolmogorov-Smirnov test (KS test) was conducted to verify the normality of the extracted voxel values from the averaged PET, fMRI and EEG measures. To determine the association between the SUV of the FDG-PET and the RA of mGluR5 and GABA_A,_ Pearson linear correlation coefficients (r) were computed with a significance level of 1% using all voxel values extracted from the averaged PET measures in GM and the core RSNs. Further, in order to elucidate the relationship between the PET measures and RS-fMRI and source-localised microstates, Pearson linear correlation coefficients (r) were computed with a significance level of 1% using averaged measures from each study. In all of the Pearson correlation analyses, the family-wise error rate (FWER), due to multiple comparisons, was controlled for using a permutation test(Groppe, Urbach and Kutas, 2011). 10^5^ Permutations were performed for each comparison (correlation), and the p-value was adjusted using the “max statistics” method(Groppe, Urbach and Kutas, 2011).

## Acknowledgements

We thank Dr. Jorge Arrubla for his assistance in FDG trimodal human data acquisition. We are grateful to Dr. Andreas Matusch for his guidance with metabolite correction for PET imaging. We are also thankful to Margo Kersey and Claire Rick for proofreading the manuscript and to Andrea Muren, Cornelia Frey, Silke Frensch and Suzanne Schaden for their technical assistance. The study was partly supported by the EU FP7-funded project TRIMAGE (Nr. 602621).

## Supplementary Figure

**Supplementary Fig. 1:**
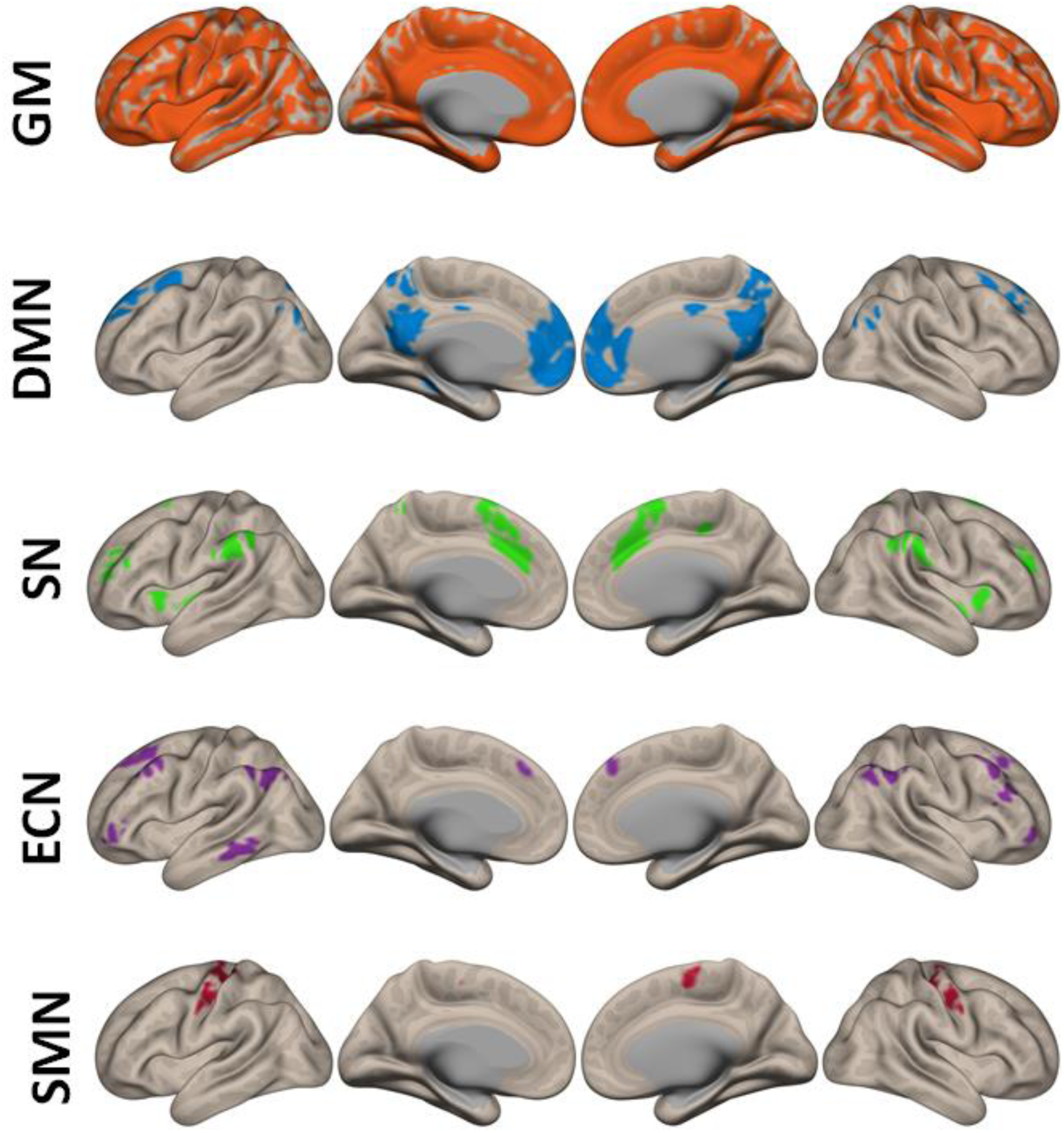
Masks of whole brain grey matter (GM) and resting state networks (default mode network (DMN), salience network (SN), executive control network (ECN) and sensory motor network (SMN)) obtained from an atlas of 90 functional regions of interest (fROIs) (Shirer *et al*., 2012).

## Supplementary Tables

**Supplementary Table 1.** Pearson Correlation r-value between averaged PET and microstates.

|  | **mGLUR5** | **GABA<sub>A</sub>** | **FDG-SUV** |
| --- | --- | --- | --- |
| **Microstate A** | 0.03** | 0.24** | 0.12** |
| **Microstate B** | 0.05** | 0.23** | 0.13** |
| **Microstate C** | 0.02** | 0.18** | 0.13** |
| **Microstate D** | 0.06** | 0.23** | 0.12** |

\*p < 0.01

**Supplementary Table 2.** Pearson Correlation r-value between averaged PET and fMRI measures in the GM and core RSNs regions.

|  | **fMRI DC** | **fMRI ReHo** | **fMRI fALFF** |
| --- | --- | --- | --- |
| **FDG SUV in GM** | 0.49* | 0.57* | 0.42* |
| **GABA<sub>A</sub> in GM** | 0.51* | 0.59* | 0.43* |
| **mGLUR5 in GM** | 0.36* | 0.38* | 0.41* |
| **FDG SUV in DMN** | 0.23* | 0.52* | 0.26* |
| **GABA<sub>A</sub> in DMN** | 0.43* | 0.62* | 0.44* |
| **mGLUR5 in DMN** | 0.10* | 0.32* | 0.30* |
| **FDG SUV in SN** | 0.27* | 0.28* | 0.24* |
| **GABA<sub>A</sub> in SN** | 0.29* | 0.31* | 0.21* |
| **mGLUR5 in SN** | 0.21* | 0.40* | 0.17* |
| **FDG SUV in ECN** | 0.43* | 0.41* | 0.35* |
| **GABA<sub>A</sub> in ECN** | 0.64* | 0.66* | 0.65* |
| **mGLUR5 in ECN** | 0.42* | 0.53* | 0.52* |

\*p < 0.01

